# Determining the Mutation Bias of Favipiravir in Influenza Using Next-generation Sequencing

**DOI:** 10.1101/375378

**Authors:** Daniel H. Goldhill, Pinky Langat, Hongyao Xie, Monica Galiano, Shahjahan Miah, Paul Kellam, Maria Zambon, Angie Lackenby, Wendy Barclay

## Abstract

**Abstract:** Favipiravir is a broad spectrum antiviral drug that may be used to treat influenza. Previous research has identified that favipiravir likely acts as a mutagen but the precise mutation bias that favipiravir induces in influenza virus RNAs has not been described. Here, we use next-generation sequencing (NGS) with barcoding of individual RNA molecules to accurately and quantitatively detect favipiravir-induced mutations and to sample orders of magnitude more mutations than would be possible through Sanger sequencing. We demonstrate that favipiravir causes mutations and show that favipiravir primarily acts as a guanine analogue and secondarily as an adenine analogue resulting in the accumulation of transition mutations. We also use a standard NGS pipeline to show that the mutagenic effect of favipiravir can be measured by whole genome sequencing of virus.

**Importance:** New antiviral drugs are needed as a first line of defence in the event of a novel influenza pandemic. Favipiravir is a broad-spectrum antiviral which is effective against influenza. The exact mechanism of how favipiravir works to inhibit influenza is still unclear. We used next-generation sequencing (NGS) to demonstrate that favipiravir causes mutations in influenza RNA. The greater depth of NGS sequence information over traditional sequencing methods allowed us to precisely determine the bias of particular mutations caused by favipiravir. NGS can also be used in a standard diagnostic pipeline to show that favipiravir is acting on the virus by revealing the mutation bias pattern typical to the drug. Our work will aid in testing whether viruses are resistant to favipiravir and may help demonstrate the effect of
favipiravir on viruses in a clinical setting. This will be important if favipiravir is used during a future influenza pandemic.

## Introduction

Influenza virus is responsible for the deaths of between 290,000-650,000 people globally each year^1^. The emergence of a novel strain of influenza in humans could lead to an influenza pandemic with significant mortality worldwide^2^. Whilst vaccination provides good levels of protection against seasonal influenza, at the start of a pandemic, antiviral drugs would be the frontline of defence during a period of development of a specific vaccine^3^. Historically, there have been only two licensed classes of antiviral drug for influenza: adamantanes and Neuraminidase inhibitors (NAIs). Adamantanes are no longer in clinical use as almost all circulating viruses are resistant^4,5^. Furthermore, some previous seasonal viruses have shown high levels of resistance to the most commonly administered NAI, oseltamivir^6^ and oseltamivir resistant A(H7N9) viruses with pandemic potential have emerged and are transmissible between ferrets^7-9^. New drugs are needed for treatment of seasonal influenza as well as for pandemic preparedness and a number of drug classes are under development including compounds that target the viral RNA dependent RNA polymerase (RdRP)^10^. In 2014, Favipiravir, an antiviral drug developed by Toyama, was licensed for use in Japan against emerging influenza viruses that exhibit resistance to other antivirals^11^. However, the exact mechanism through which favipiravir exerts an antiviral effect on influenza is unclear. An increased knowledge of the mechanism of action of favipiravir could be useful in determining whether specific viruses are less susceptible and evaluating the potential for emergence and transmission of resistant viruses.

Favipiravir is a nucleoside analogue that is active against all subtypes of influenza and has shown a potent antiviral effect both *in vitro* and *in vivo^12-17^.* Favipiravir has completed a phase III clinical trial in Japan and has undergone a phase III trial in the USA^18^.

Favipiravir has also been shown to be active *in vitro* and in animal models against a wide range of RNA viruses, some for which there are no licensed drugs as a treatment option^18-25^. There is strong evidence that favipiravir acts as a mutagen by incorporating into both positive and negative stranded RNA and being aberrantly copied as multiple bases^15,26-30^.

This is thought to be a different mechanism of action from ribavirin, another broadly acting nucleoside analogue that has been used previously to treat influenza^26,31^. Studies have shown that favipiravir competes against guanine and adenine to be incorporated into RNA and is non-competitive against cytosine and uracil^30,32-34^. This would suggest that favipiravir acts as a purine analogue and should cause mostly transition mutations. Studies measuring the mutation bias of favipiravir in influenza have had mixed results. Baranovich *et al.* used Sanger sequencing of virus passaged in presence of drug to show a C->U and a G->A mutation bias as expected but also saw a G->U mutation bias after 48hrs of exposure to favipiravir^27^. Vanderlinden *et al.* also used Sanger sequencing to show a C->U and G->A bias following a passaging experiment and showed an increase in Shannon entropy using next-generation sequencing (NGS)^35^. However, in contrast to studies using Sanger sequencing, Marathe *et al.* reported a slight bias towards transversions in influenza infected mice treated with favipiravir using NGS^36^. Studies with other viruses have given mutation patterns which suggest that favipiravir acts as a purine analogue^28,29,37,38^. Interestingly, several studies with favipiravir and influenza have suggested that favipiravir acts not as a mutagen but as a chain terminator preventing the extension of the RNA strand following incorporation^32,33^. A primer extension study suggested that the block could occur with a single molecule of favipiravir^32^ but other studies have suggested that chain termination occurs following the incorporation of two molecules of favipiravir^30,33,34^.

In this study, we used next generation sequencing to determine the mutation bias of favipiravir on influenza virus RNAs. We employed two methods of analysis: the first method uses Primer ID which is a technique for labelling each individual RNA molecule with a barcode to account for PCR and sequencing errors^39-41^. This technique can very precisely uncover the mutation bias by analysing small, targeted areas of the genome. The second method developed a novel analysis of data obtained from a standard sequencing pipeline as would be found in many National Influenza Centres or public health laboratories. This showed the mutation bias induced by drug treatment over the whole genome was similar to that detected using the precise Primer ID methodology and confirmed that the effect of favipiravir could be readily measured using NGS from a standard sequencing pipeline.

## Methods

### Reagents, Cells and Viruses

Favipiravir was kindly provided by Toyama Chemical Company under an MTA and reconstituted in DMSO and frozen into aliquots. MDCK and 293-T cells were grown in Dulbecco’s modified Eagle’s medium (DMEM; Gibco) with the addition of 10% Fetal Bovine Serum (labtech.com), 1% non-essential amino acids (Gibco) and 1% penicillin/streptomycin (Sigma-Aldrich). A/England/195/2009 (Eng195) is an early isolate from the 2009 A(H1N1) pandemic provided by Public Health England (PHE).

### Minigenome Assay

Four pCAGGS plasmids encoding the polymerase (PA, PB1 and PB2) and NP from influenza A/England/195/2009 A(H1N1)pdm09 virus, were transfected using Lipofectamine 3000 (Invitrogen) into 293T cells in 24 well plates. In addition, we transfected plasmids directing expression from a PolI promoter of either a Firefly luciferase gene in negative sense flanked with influenza A non-coding sequence from the NS segment or the HA gene segment from influenza A/Victoria/3/75 H3N2 virus (Vic75), and a PolII Renilla luciferase plasmid as a transfection control. Cells were lysed with 200μl of passive lysis buffer (Promega) and polymerase activity was measured using Dual-Luciferase Reporter Assay (Promega) on the FLUOstar Omega plate reader (BMG Labtech). Polymerase activity is reported as Firefly luciferase activity normalized by Renilla activity.

### Next-Generation Sequencing with Primer ID

At 24 hours after transfection, 293T cells from the minigenome assay were lysed and RNA was extracted using the RNA mini kit (Qiagen). The reverse transcription primer for primer ID (5’-TGCGTTGATACCACTG CTTTNNNNTNNNNTNNNN CCCAGTCCAAGTGAAACCCT C-3’) consisted of a PCR tag, random barcode of the form NNNNTNNNNTNNNN and sequence specific to the H3 HA. Reverse transcription was performed with Superscript III (Thermo Fisher). qPCR using SYBR green (Thermo Fisher) was used to calculate the number of cDNA molecules to use for each PCR reaction. 20,000-40,000 molecules were used for each reaction. The PCR primers were 5’-CGGGGAAAATATGCAACAATCCT-3’ and 5’-TGCGTTGATACCACTGClll. The PCR product was designed to be 279 bases to avoid any fragmentation step during sample preparation ensuring the barcode was not sheared from the sample. Sample preparation was performed using the NEBNext Ultra kit (NEB). Samples were sequenced giving 150bp paired end reads on an Illumina MiSeq. Sequencing data for the samples were processed and analysed using custom scripts in Python and R. Reads were first paired to form a single sequence and subjected to quality control using QUASR v7.01^42^ to retain reads with a median phred score of 20 and minimum read length of 250bp. Intact barcode sequences were extracted from the read pairs; any sequences without a fully formed barcode or with errors in the internal Ts of the barcode were discarded. Consensus sequences were generated for each barcode that had more than three reads with the consensus taken as the majority of the reads. Samples for which there was no majority read were discarded as potentially this could be an example of two RNA sequences having the same barcode^43^. The consensus sequences were mapped and compared to the Vic75 reference and any variants were extracted. We subsequently decided to use a more stringent cut-off of four reads per barcode to minimize errors caused by barcodes with a low number of reads. We present all our sequencing results as mutations in positive orientation as would have been seen in the mRNA.

### qPCR

RNA was extracted from the mini-genome assay. Specific primers were used to reverse transcribe mRNA from the firefly luciferase as previously described^44^. qPCR was performed with SYBR Green using 18S RNA as a control. ΔΔCt was calculated and the results are shown normalized to the drug free control.

### Next-Generation Viral Sequencing with Primer ID

1.2*10^6 cells were inoculated with Eng195 at a MOI of 1.5 and incubated at 37°C for 18 hours in serum free media with added 1μg/ml trypsin (Worthington) and with different concentrations of favipiravir diluted in DMSO. Control wells contained DMSO without favipiravir. After 18 hours, samples were taken from the supernatant and plaqued on MDCK cells to determine final viral titre. RNA was extracted from the cells using RNEasy kit (Qiagen). Sequencing was performed as described above with the exception that the Primer ID RT primer contained sequence specific for PB1 vRNA (5’-TGTCCAGCACGCTTCAGGCTNNNNTNNNNTNNNNAGAAGATGGTCACGCAAAGAA-3’) and the PCR product was 302 bases long including the PCR primers (’-TCACAACATTTGCCAGTTTGG-3’,-TGTCCAGCACGCTTCAGGCT-3’). On analysing the sequencing data, a site which varied considerably in all samples was detected which was likely a polymorphism in the initial population. This site was removed from all analyses.

### Next-Generation Sequencing without primer ID

1.2*10^6 cells were inoculated with England 195 at a MOI of 1 and incubated at 37°C for 24 hours as described above. Control wells contained DMSO but no favipiravir. After 24 hours, samples were taken from the media and titred on MDCK cells by plaque assay. Whole genome next generation sequencing was performed using a pipeline at Public Health England. RNA was extracted from viral lysate using easyMAG (bioMérieux). One step Reverse-Transcription-PCR was performed with Superscript III (Invitrogen), Platinum Taq HiFi Polymerase (Thermo Fisher) and influenza specific primers^45^. Samples were prepared for NGS using Nextera library preparation kit (Illumina). Samples were sequenced on an Illumina MiSeq generating a 150-bp paired end reads. Reads were mapped with BWA v0.7.5 and converted to BAM files using SAMTools (1.1.2). Variants were called using QuasiBAM, an in-house script at Public Health England. Samples were compared using a permutation analysis to calculate the probability of a magnitude of mutation bias as great as observed given the mutations in the samples. Permutation analyses were performed in R with 10,000 iterations for each analysis. Mutations were randomised between two samples maintaining the number of mutations found within each sample. The magnitude of the mutation bias was then calculated as the sum of the absolute value of the difference in the relative proportions of each mutation type. The p value was then calculated as the number of iterations/10000 with a value greater than the observed value. A further permutation analysis calculated the probability of a bias of guanine analogue mutations (e.g. C->U and G->A). This analysis was performed as above but only used the sum of the absolute value of the difference in the relative proportions of C->U and G->A.

## Results

### Primer ID allows calculation of mutation bias and relative mutation rate

In order to determine the mutagenic effect of favipiravir, we employed next generation sequencing using Primer ID to analyse the products of a minigenome assay^46^, which allowed for the unbiased measurement of mutations (Figure 1). When sequencing virus, particularly over several rounds of replication, a proportion of possible mutations will not be measured as they would cause too large a fitness cost to the virus and thus will not be amplified. To avoid this scenario, we sequenced the reporter gene from the minigenome assay as the reporter protein has no effect on further RNA accumulation. Thus, this strategy should reveal the complete spectrum of mutations caused by replication in the presence of favipiravir. Primer ID is a method which labels each molecule of RNA with a unique barcode (Figure 1). This method allowed us to examine a large number of independent mutational events as each mutation could be associated with an individual RNA molecule. In addition, by comparing multiple sequencing reads with the same barcode, we could remove sequencing errors as these would not appear in the majority of the reads. The sample without favipiravir provides a baseline mutation rate consisting of the background mutation rate of the influenza virus polymerase plus mutations caused by the reverse transcriptase during reverse transcription. Drug-treated samples can be compared to this sample to measure how favipiravir increased the mutation rate.

**Figure 1.**
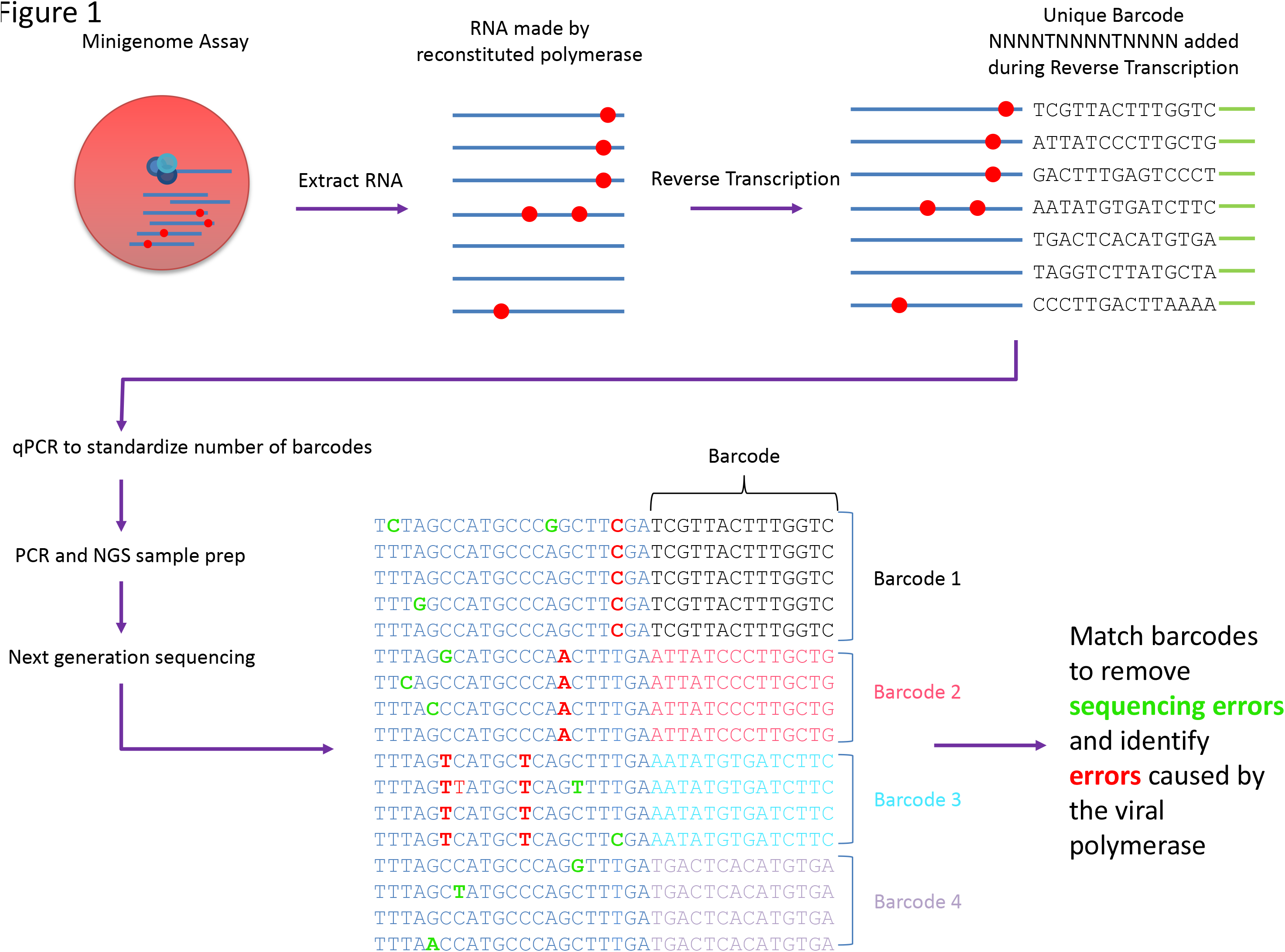
Primer ID Method for determining mutation bias. RNA was extracted and a unique barcode of the form NNNNTNNNNTNNNN added during reverse transcription. qPCR was used to standardize the number of barcodes for NGS. Samples were sequenced and barcodes matched to allow removal of PCR and sequencing errors.

We reconstituted influenza RdRP in situ by expressing the polymerase proteins and nucleoprotein from transfected plasmids. We introduced two viral-like RNA templates, one in which the authentic open reading frame was replaced by the firefly luciferase gene, and one that represented RNA segment 4 and encoded H3 haemagglutinin (HA). The transfected cells were incubated in the presence of favipiravir. Increasing concentrations of favipiravir from 1 to 100 μM caused a reduction in the activity of the luciferase reporter (Figure 2A). However, qRT-PCR analysis of the amount of H3 HA mRNA accumulated revealed no decrease in mRNA levels that would account for the loss of luciferase activity at least up to 50 μM drug (Figure 2B). At 100 μM favipiravir, there was a significant reduction in mRNA (P<0.0001). This suggested that at doses up to 50 μM, the inhibitory effect of favipiravir in the minigenome assay was caused by mutagenesis and not through chain termination, which could have played a role at the highest dose of drug.

**Figure 2.**
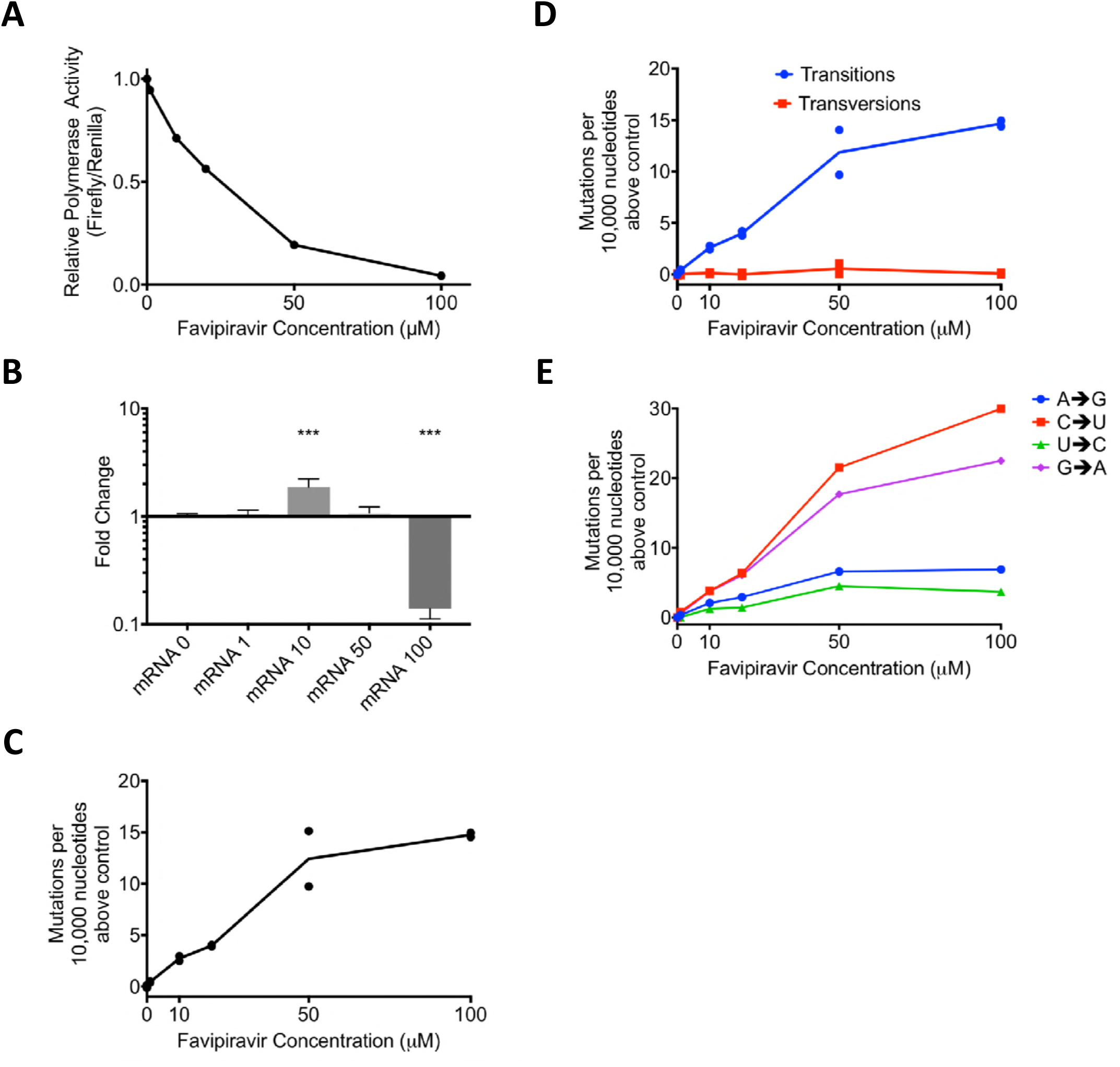
Favipiravir causes transition mutations which reduces polymerase activity in a minigenome assay. **A)** Minigenome Assay. Plasmids were transfected into 293-T cells and favipiravir was added. At ~21hrs the cells were lysed and luciferase activity was measured. The relative polymerase activity is calculated as firefly activity / renilla activity. **B)** A reporter plasmid (HA pol1) from the minigenome above was sequenced using Primer ID and NGS. The mutations were tallied as described in Methods. Two independent biological samples were sequenced for each concentration of drug in the same sequencing reaction. The number of mutations per 10,000 nucleotides above the average of the two control samples were compared for each sample. **C)** The number of mutations per 10,000 nucleotides above the control for each sample was calculated for transitions and transversions. **D)** As 1C calculated for each class of transition mutation. The values are calculated as the mutation rate for an individual base. The average for the two samples is plotted. **E)** qPCR was performed on the luciferase reporter mRNA from a minigenome assay. ΔΔCt was calculated using 18sRNA and results are shown normalized to the drug-free control. N=6. *** p<0.001.

In order to test how favipiravir affected the mutation rate of the reconstituted viral polymerase, we sequenced the positive stranded H3 HA RNAs. As each individual barcode represents a single RNA molecule, we calculated consensus sequences for each barcode. Mutations which did not appear in a majority of reads were ascribed to PCR or sequencing error and removed from further analyses. In total, we analysed 6,623 substitutions in ~6,900,000 bases of sequencing data. Figure 2C shows the number of mutations per 10,000 nucleotides above the baseline (0 μM favipiravir) for each sample. As the concentration of favipiravir increased, the number of mutations increased. At the highest concentration of favipiravir tested (100 μM), there would be an additional 15 errors per 10,000 nucleotides on average compared to the control. We varied the cut-off for the number of sequencing reads needed to include a barcode (Supplemental figure 1). The choice of cut-off did not significantly alter the results for values <10 reads. We chose a cut-off of 4 reads per barcode as this removed some errors associated with low numbers of reads per barcode whilst including the majority of the data.

We next categorised the mutations identified by sequencing as transitions or transversions, or as the individual base-pair mutations (Figure 2D, E). Our results confirmed that the main cause of the increase in mutation rate was transition mutations (Figure 2D). There was no increase in the rate of transversion mutations as the concentration of favipiravir increased (F-test, F= 0.4593, d.f. 1,4, p=0.5351). Figure 2E shows the increase in the likelihood of different categories of mutations compared to the control. The most common transitions were C->U and G->A mutations that would be induced when favipiravir is acting as a guanine analogue. However, there was also a smaller increase in the reverse transitions from U->C and A->G where favipiravir acts as an adenine analogue. On average, there was an approximately 3.5-fold increase in the rate of C->U or G->A mutations compared to a U->C or A->G mutations.

### Primer ID sequencing of viruses confirms that favipiravir causes mutations

We next tested whether we could use Primer ID to measure the increase in mutation rate of RNAs generated during virus infection. To minimize the loss of viral RNAs that contained mutations rendering the virus nonviable, we infected cells at a high MOI so that there was only a single replication cycle. We first confirmed that favipiravir inhibited influenza under these conditions (Figure 3A). There was a greater than 1000-fold reduction in infectious titre of influenza A/Eng195/2009 A(H1N1)pdm09 virus (Eng 195) after 24 hours infection at high concentrations of favipiravir and a 10-fold reduction at 1 μM drug. We extracted RNA from the cells and sequenced the vRNA of RNA segment 2 with appropriate barcoded primers. In total, we analysed ~56,000,000 bases and found 25,441 substitutions. All concentrations of favipiravir showed an increase in mutation rate compared to the no drug control (Figure 3B). The mutation rate caused by favipiravir was ~3 fold higher at 10 μM than at 1 μM, but surprisingly, the mutation rate at 100 μM favipiravir was lower than at 10 μM. The increase in mutation rate at all concentrations of favipiravir was almost entirely due to transitions (Figure 3C). The mutation bias measured was subtly different than that seen using the minigenome assay with C->U occurring most often but G->A and U->C mutations occurring at comparable rates (Figure 3D). This suggests that there was a higher rate of incorporation of favipiravir during negative strand synthesis compared to positive strand synthesis in virus infected cells (see Figure 5).

**Figure 3.**
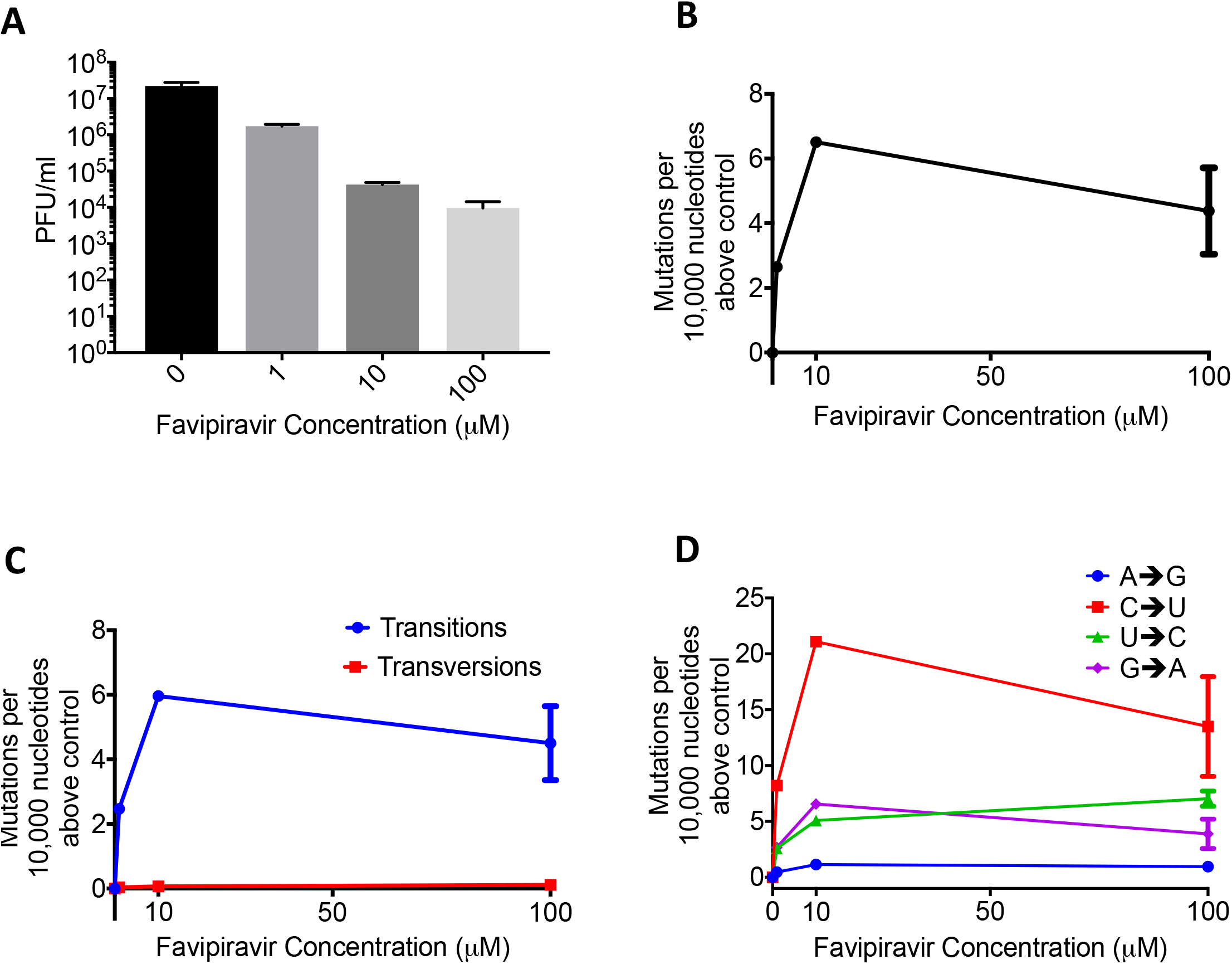
Favipiravir causes transition mutations reducing viral fitness. **A)** Virus was added to MDCK cells at a high MOI of 1.3 and favipiravir was added at an appropriate concentration diluted in DMSO. The supernatant was plaqued after 20 hours and the titre calculated in plaque forming units/ml. N=3. **B)** After 18 hours the cells were lysed and the RNA extracted for sequencing using Primer ID. The number of mutations per 10,000 nucleotides above the control was plotted for each sample. **C)** The number of mutations per 10,000 nucleotides above the control for each sample was calculated for transitions and transversions. **D)** The number of mutations per 10,000 nucleotides above the control for each sample was calculated for each class of transition mutation. The values are calculated as the mutation rate for an individual base.

**figure 5.**
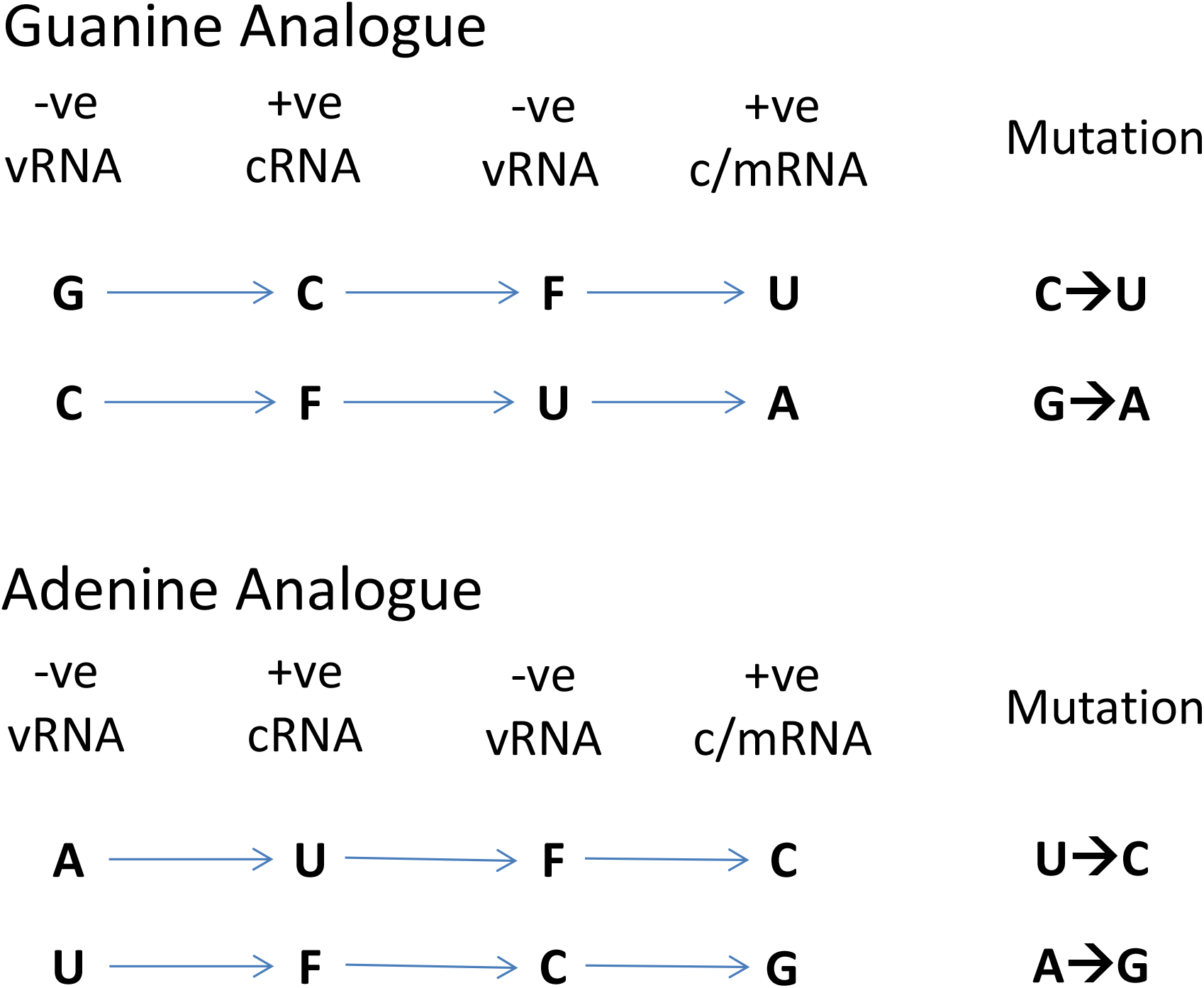
A schematic showing how favipiravir causes mutations during +ve and -ve strand synthesis.

### Next generation sequencing can reveal mutation bias

The experiments with Primer ID showed the mutation rate and bias for a small targeted portion of influenza genome. Next, we wanted to test whether we could measure the mutagenic effect of favipiravir using a standard NGS pipeline typical of those in public health laboratories (Supplemental figure 2). Eng195 virus was propagated at a high MOI for 24 hours in the presence of 10 or 100 μM favipiravir. The supernatant was plaqued to confirm that favipiravir had an inhibitory effect on the virus and there was >2 log inhibition at 10 μM and >4 log inhibition at 100 μM. We extracted RNA from virus particles in the supernatant and used next generation sequencing to obtain sequence data from the population of surviving viruses. In order to analyse mutation bias using next generation data, it is necessary to ensure that the mutations used for the analysis are independent so that the same mutation occurring on multiple reads is not counted as multiple mutational events but as a single mutational event. Therefore, we treated each base in the influenza genome independently and recorded only the most common mutation (if any) for each site (Supplemental figure 2). Taking these sites in aggregate will give a combination of true mutations as well as other sources of error, most notably sequencing error. Figure 4 shows the sum of mutations over the whole genome for viruses propagated in 10 μM or 100 μM favipiravir or for control viruses which were not exposed to favipiravir. Comparing the pattern of mutations between the control viruses and the viruses exposed to drug allowed us to control for sequencing errors. The pattern of mutations seen in both samples exposed to favipiravir were significantly different to the control (Permutation analysis, p<1*10^-4; Supplemental figure 3A, 3C.) The mutation bias was caused by an excess of C->U and G->A transitions compared with the control viruses (Permutation analysis, p<1*10^-4, Supplemental figure 3B, 3D). There was no significant difference between the mutation bias at the two different concentrations of favipiravir tested (Permutation Analysis, p=0.26, Supplemental figure 3E, 3F). To demonstrate further that this method measures a true mutational signal, we took the 500 sites with the highest degree of polymorphism and repeated the analysis (Supplemental figure 4). The new analysis showed an increased effect size strongly suggesting that mutations caused by favipiravir lead to a signal in the sequencing data that is not masked by sequencing error. We chose to use the relative proportion of the mutation types to compare between samples as opposed to the absolute number of polymorphisms. This was a conservative choice as there may be biases between samples which could affect the absolute number of polymorphism due to the number of viruses in the sample.

**Figure 4.**
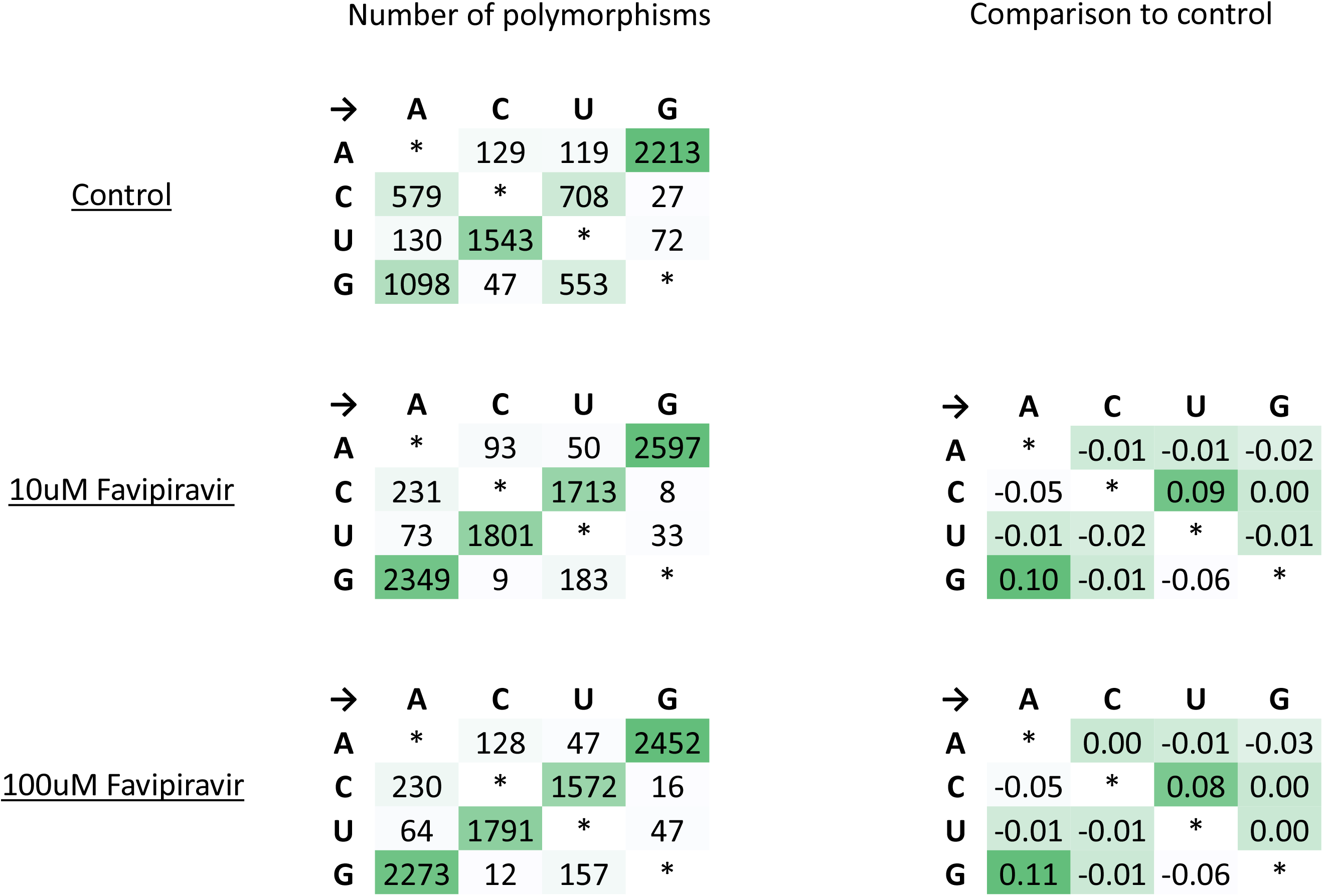
Next-generation sequencing data shows that favipiravir acts as a guanine analogue. Virus was added to MDCK cells at a high MOI of 1 and drug was added as previously described. Supernatant was taken and was sequenced and analysed as described in *Methods.* The most common polymorphism for each base is shown for virus exposed to drug and to a drug free control. The comparison shows the difference in percentage for each class of mutations revealing mutation bias.

## Discussion

In this study, we used two different methods of analysing next-generation sequencing data in order to show that favipiravir acts as a mutagen with a distinct bias to induce transitions in influenza virus RNAs. The first method used Primer ID to measure precisely the increase in mutation rate and the mutation bias of the influenza polymerase caused by favipiravir in an *in vitro* system. We confirmed that favipiravir has a bias for transition mutations and acts as a purine analogue^17,26,32,33^. We were able to demonstrate that favipiravir competed primarily with guanine and secondarily with adenine resulting in an increase in C->U and G->A mutations at higher concentrations of drug and a lower rate of increase in U->C and A->G mutations (Figure 5). The second method used data from whole-genome sequencing of viruses that had been exposed to favipiravir during single cycle replication and showed that viral populations exposed to favipiravir had a distinct bias for transition mutations, specifically C->U and G->A mutations.

Previous methods of sequence analysis for determining mutation bias in influenza RNAs induced by favipiravir have relied on Sanger sequencing of individual viral clones^27,31^. This technique is laborious and results in the detection of relatively few mutations: on the order of 100 mutations for an entire experiment^27,31^. Furthermore, the technique can be biased due to selection of beneficial mutations which may appear in multiple clones or to accidentally counting an initial polymorphism in the population as a mutational event that occurred in multiple clones. Sequencing a small region of the genome across many clones is especially prone to this error. Next-generation sequencing with Primer ID is a powerful technique which allowed us to examine orders of magnitude more mutations than Sanger sequencing and was less prone to biases present in examining a small number of mutations. Primer ID allowed us to remove sequencing error from next-generation sequencing data and to detect changes in mutation rate and mutation bias^39,40^. Primer ID identified thousands of mutations in a single sample exposed to favipiravir, a number which would be impractical using Sanger sequencing. We were able to show that favipiravir acts as both a guanine and an adenine analogue whereas Sanger sequencing was not sensitive enough to measure the lower rate of adenine mutations^27^.

The use of the minigenome assay allowed us to see all mutations generated by polymerase and not just those that would allow viable viruses. Pauly *et al.* have recently shown that the mutation rate for influenza has been significantly underestimated by only counting mutations which occur in plaque forming viruses^47^. Sequencing only viruses which have exited the cell ignores mutations that cause defects in packaging or cellular exit. By contrast, as the mRNA from the reporter in the minigenome assay is not translated to a protein that can impact on viral fitness, the full spectrum of drug-induced mutations can be seen. Allowing for multiple rounds of virus replication makes it difficult to see strongly deleterious mutations, which make up a significant proportion of the mutations for influenza, because they are selected against^48^. The minigenome assay has no selection on mutations and does not suffer from this bias. However, when we used a Primer ID approach to sequence a small portion of the viral genome from PB1 amplified during virus infection rather than in the minigenome assay, we found, contrary to the minigenome sequencing, that there was no increase in the mutation rate at the highest concentrations of favipiravir. This is likely due to selection against deleterious mutations that occurs even in a single cycle of replication.

Favipiravir causes mutations randomly and therefore there will be a distribution in the number of mutations during each strand replication. Some RNAs will have many mutations whereas others will have fewer. The majority of the RNA that was sequenced will come from viruses which have suffered few mutations, as viral RNAs with more mutations will interfere with ongoing replication. Therefore, the more successful favipiravir is at causing mutations, the greater the bias to sequencing the small number of viruses with fewer mutations. This most likely explains why the mutation rate we measured appeared lower at 100 μM favipiravir than at 10 μM.

Although Primer ID can remove sequencing error, it is still impossible to distinguish between errors due to the flu polymerase and the reverse transcriptase used during the Primer ID reaction. A recent paper has suggested that care must be taken as these two error rates are the same order of magnitude^47^. For this reason, we have not reported an absolute error rate but a relative error rate compared to the drug-free baseline sample. However, for our experiments, the mutation rate caused by favipiravir was much higher than the calculated baseline mutation rate caused by reverse transcription errors plus errors naturally caused by the influenza polymerase. Furthermore, as all samples underwent identical processing, there is no reason to believe that the error rate during reverse transcription differed between samples and therefore, this is unlikely to bias our data. Care would need to be taken before comparing samples which have not been prepared concurrently especially if using different reverse transcription enzymes.

One disadvantage to Primer ID is that it sequences only a small part of the genome. This potentially could lead to mutation biases if that part of the genome was under strong selection or due to local sequence structure. As we sampled only one region of the HA, we could not test whether there were specific structural differences between the HA sequence and other flu segments leading to mutational hotspots. However, the similarity between our analysis of RNAs from primer ID vs whole genome sequencing suggests we did not inadvertently sample a mutational hotspot. The precision and ease with which Primer ID was able to distinguish mutation bias and observe changes in mutation rate leads us to suggest that it could become a standard method for analysing the effects of nucleoside analogues and other mutagenic drugs.

Our second method of analysis sequenced the whole flu genome in populations of viruses that had been exposed to favipiravir and a control population that was not exposed to the drug as might be found in a clinical setting. The main disadvantage of this technique is that it is unable to distinguish between sequencing error and ‘true’ errors caused by the flu polymerase. Therefore, it is not possible to quantify the actual number of errors due to polymerase nor was the method sensitive enough to demonstrate any increase in the rate of U->C and A->G mutations. Despite these limitations, there are several advantages to this method that may prove to be of use in clinical settings. This method is extremely simple to use as the viruses can be entered into the standard influenza sequencing pipeline without any additional processing steps and could also be used to reanalyse data that had been previously collected. The analysis also encompasses the whole genome and so is resistant to any biases caused by local RNA structure nor is it biased by single polymorphisms that may have been present in the initial populations. If favipiravir is used in a clinical setting, this method may be a simple way to show that favipiravir is having a measurable effect by comparing viral mutations in pre-treatment and post-treatment samples.

In contrast to our finding that favipiravir acts as a purine analogue, a previous study that used NGS to determine the mutation bias of favipiravir *in vivo* found an excess of transversion mutations^36^. The analysis in Marathe *et al.* counted each individual NGS read as a separate mutational event, which may have led to a bias, as mutations from pre-existing polymorphisms or mutations that are positively selected will be counted multiple times. By contrast, our method of analysing NGS data ensured that mutations were independent by only counting one mutation at each site in the genome (Supplemental figure 2). Many recent papers that analyse NGS data use a cut off e.g. 5% or 1% of reads below which variants are not counted^31,36,38^. However, using a cut-off discards a large amount of the sequence data as only a small proportion of sites are included. Our analysis (Figure 4, Supplemental figure 2) used all the sequencing data without imposing a cut-off and this led to increased noise in the data but ensured that there was no bias towards pre-existing polymorphisms or variations in sequencing depth. We also tested the mutational bias by only counting the 500 sites with the largest degree of polymorphism (Supplemental figure 4) which showed similar results to our main analysis though potentially with less noise. This suggests that imposing a cut-off on variants will not bias the results if the sequencing contains enough variants that positive selection and pre-existing polymorphisms are unlikely to influence the results.

Our data showed that favipiravir acts as a mutagen with a bias towards transitions in agreement with most other studies of this drug’s effect on RNA viruses^27,28,35^. We found that at lower concentrations of favipiravir, there was no evidence that the drug was acting as a chain terminator as there was no reduction in the amount of mRNA despite a reduction in reporter gene activity (Figure 2A, B). At the highest concentration tested (100 μM), there was a reduction in mRNA which could have been caused by chain termination or through introduced mutations preventing RNA replication. The lack of evidence for chain termination at lower concentrations of favipiravir suggests that favipiravir is primarily acting as a mutagen. Biochemically, favipiravir acts as a purine analogue binding to either C or U in place of G or A respectively. The most common mutations caused by favipiravir were C->U and G->A. These mutations were caused by favipiravir binding to C in place of a G on the positive or negative strand synthesis and subsequently pairing with a U in the next synthesis cycle (Figure 5). The reverse transitions caused by favipiravir binding to U happened at a ~3.5-fold lower rate. This confirms that favipiravir is most competitive against G as had been previously seen in primer extension assays^32,33^.

Next-generation Sequencing is a powerful technique for analysing mutational data and determining mutational biases. Care must be taken to perform analyses which minimize potential biases by ensuring that mutations are only counted when they occur independently of each other. We used NGS to show that favipiravir is acting as a mutagen causing multiple additional mutations per influenza genome on average at higher concentrations of favipiravir. Lethal mutagenesis of influenza is a viable antiviral strategy and may be difficult to evolve resistance against clinically^49^. Our increased knowledge of the precise mechanism of favipiravir means that we are better placed to test whether the drug is having a clinical effect as well as to see whether viruses are becoming resistant to favipiravir. This will be important when this drug is used in a pandemic situation.

## Acknowledgments

This work was supported by the National Institute for Health Research Health Protection Research Unit (NIHR HPRU) in Respiratory Infections at Imperial College in partnership with Public Health England (PHE). PL, PK and WB were supported by Wellcome Trust Grant 200187/Z/15/Z. The views expressed are those of the authors and not necessarily those of the NHS, the NIHR, the Department of Health or PHE.

---

**Supplemental figure 1.**
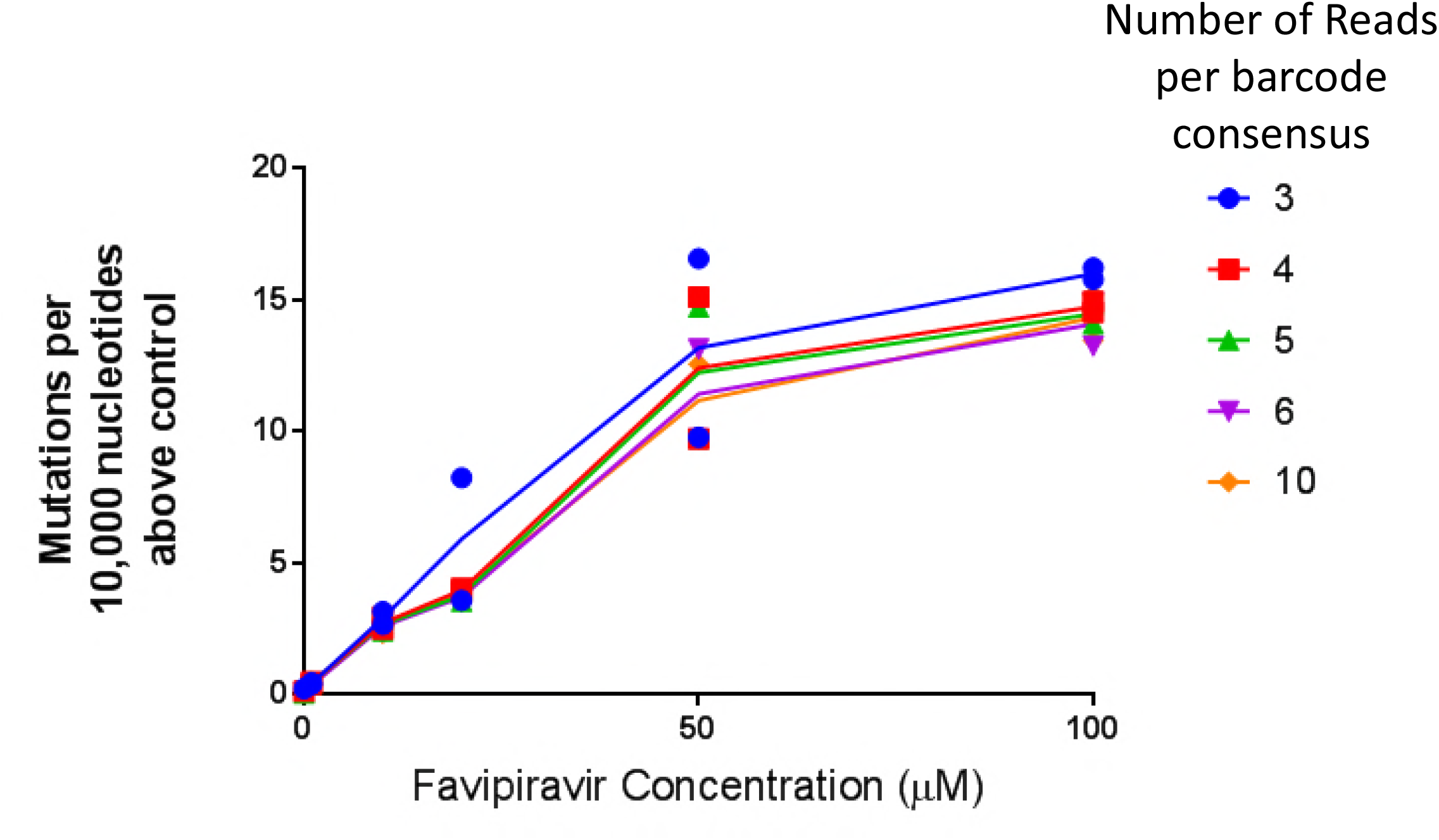
Determination of the optimal cut-off for number of reads per barcode. The cut-off for the number of reads necessary to include a barcode was systematically varied and the number of mutations per 10,000 nucleotides above the control plotted for each sample.

**Supplemental figure 2.**
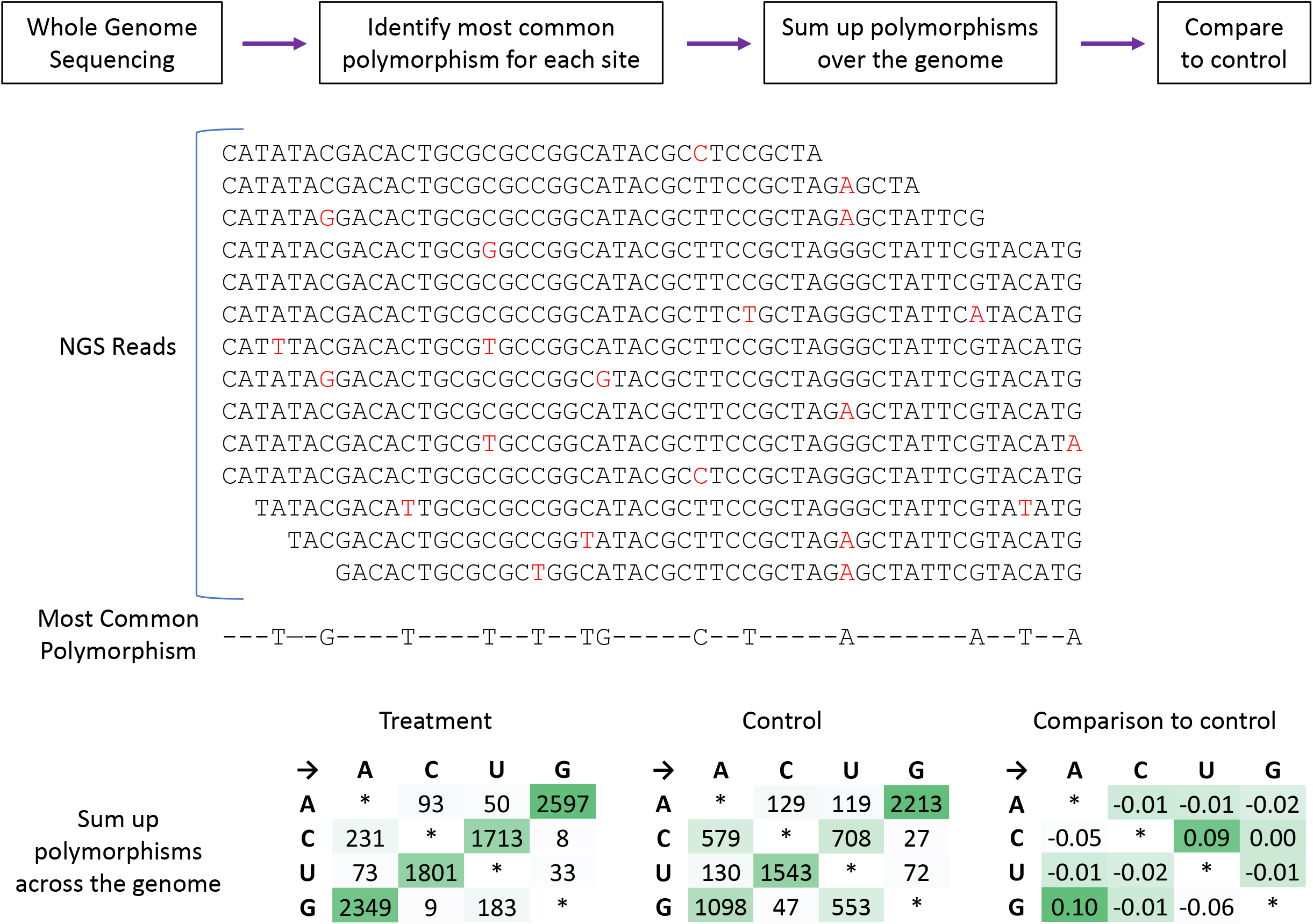
A method to analyse mutation bias from whole genome NGS data. Whole genome sequencing data from a standard pipeline was aligned to a reference. The most common polymorphism for each site in the genome was calculated. These polymorphisms were summed up and the mutation bias of different samples can be compared.

**Supplemental figure 3.**
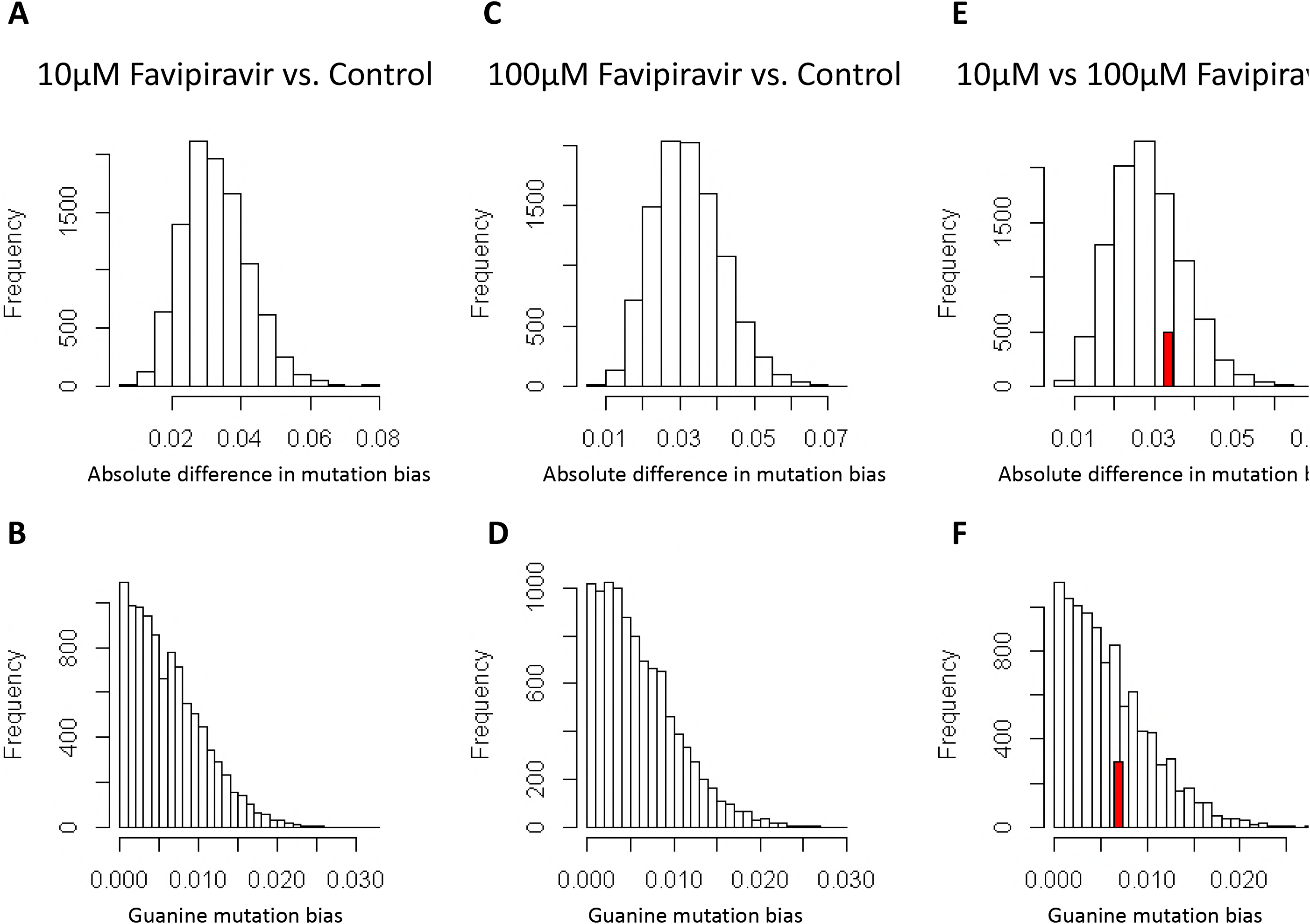
A permutation analysis was performed on the mutation data. The substitutions were randomized between the treatment and control and either the total difference in mutation bias was calculated (A, C, E) or the bias for acting as a guanine: analogue (B, D, F.) 10,000 permutations were performed for each analysis. The red bars ¦ show the observed value where it occurs within the values generated by the permutations. **A)** The mutation bias for 10 μM favipiravir was compared to the control (observed value = 0.39; <1*10^-4). **B)** The difference in bias for guanine mutations (observed value = 0.19; ι <1*10^-4). **C)** The mutation bias for 100 μM favipiravir was compared to the control (observed value = 0.37; <1*10^-4). **D)** The difference in bias for guanine mutations ¦ (observed value = 0.19; <1*10^-4). **E)** The mutation bias for 10 μM favipiravir was I compared to 100 μM favipiravir (observed value = 0.03; p=0.26). **F)** The difference in bias for I guanine mutations (observed value = 0.007; p=0.34).

**Supplemental figure 4.**
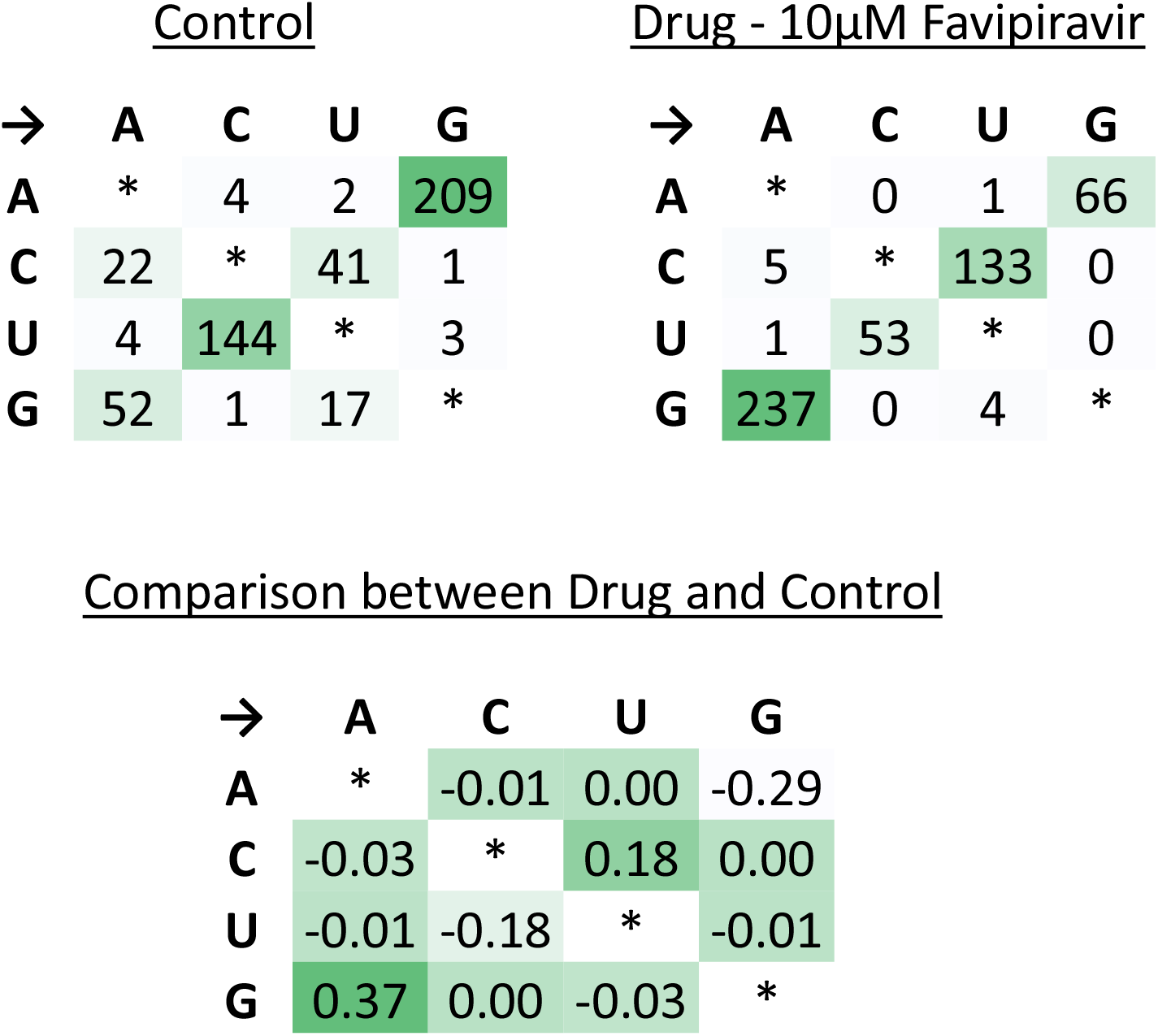
The same data from Figure 3 was reanalysed using only the 500 sites: with the largest degree of polymorphism.

